# Navigating the Maze of Mass Spectra: A Machine-Learning Guide to Identifying Diagnostic Ions in O-Glycan Analysis

**DOI:** 10.1101/2024.06.28.601175

**Authors:** James Urban, Roman Joeres, Luc Thomès, Kristina A. Thomsson, Daniel Bojar

## Abstract

Structural details of oligosaccharides, or glycans, often carry biological relevance, which is why they are typically elucidated using tandem mass spectrometry. Common approaches to distinguish isomers rely on diagnostic glycan fragments for annotating topologies or linkages. Diagnostic fragments are often only known informally among practitioners or stem from individual studies, with unclear validity or generalizability, causing annotation heterogeneity and hampering new analysts. Drawing on a curated set of 237,000 *O*-glycomics spectra, we here present a rule-based machine learning workflow to uncover quantifiably valid and generalizable diagnostic fragments. This results in fragmentation rules to robustly distinguish common *O*-glycan isomers. We envision this resource to improve glycan annotation accuracy and concomitantly make annotations more transparent and homogeneous across analysts.

## Introduction

Glycans decorate proteins and lipids and are present in all biological taxa [1]. Molecules interacting with glycans, such as lectins, are frequently sensitive to the three-dimensional conformation of a glycan [2], largely dictated by its constituent monosaccharides and the linkages joining them together. Small changes in linkage or hydroxyl group orientation can lead to different 3D structures with significant biological effects as consequences, for instance by differentially stabilizing a protein depending on the sialic acid linkage [3] or yielding qualitative differences in lectin binding depending on the exact glycan sequence [4]. This makes detailed characterization of glycan sequences in glycomics data crucial for uncovering the roles of specific isomers in particular biological systems. While many methods can be used for this purpose, we will focus our attention here on the most common approach: tandem mass spectrometry, usually preceded by liquid chromatography to separate isomeric structures.

Diagnostic fragments—only, or at least preferentially, occurring in one isomer—comprise a substantial part of current and preferred annotation strategies, due to their ease of use compared to alternative strategies such as exoglycosidase digestion. Examples here include diagnostic fragments to distinguish sialic acid linkage in *N*-glycans [5] or for the distinction of Lewis A and X structures [6]. Despite this, the usage of diagnostic fragments is neither standardized nor formalized, creating a lack of transparency and an entry barrier for analysts. No central databases or resources exist to catalog or compare such diagnostic fragments. Further, this lack of formalization also means that no quantitative confidence value can be attached to an individual human annotation, withholding necessary context and hampering transparency.

Several comprehensive studies to identify diagnostic fragmentation have been carried out before [5–8]. Typically, isomer-specific rules are devised or evaluated based on spectra obtained in a single experiment; often one carried out for the express purpose of finding these fragments and collected by the same person(s) that then analyzes it to that effect [6–8]. In rare cases, these rules are then validated on different experimental set-ups [5]. Yet, often, little information exists about whether, or to which extent, commonly used diagnostic ions are generalizable to different set-ups. Further, the quantitative efficacy of most rules is typically unknown, as well as the efficacy of combining multiple rules derived from disparate experiments, making them essentially soft rules, in which the (prominent) presence of an indicated fragment is associated with an undetermined annotation confidence.

Given the prevalent use of single/double fragment presences to determine structural details, evaluating and quantifying the performance of such criteria could not only improve annotation accuracy but also attach a confidence level to each annotation, allowing for a proper evaluation of attached biological findings. Here, we will focus on *O*-glycans as a test case. *O*-glycans are fantastically diverse in the context of mucin glycosylation [9] and very much dependent on diagnostic fragments in their annotation, due to a less rigid biosynthesis than *N*-glycans. Recent comparisons across different analysts in the area of *O*-glycoproteomics have highlighted substantial heterogeneity [10] and it is to be expected that a similar situation arises in *O*-glycomics, especially for new analysts, due to the lack of resources and challenging nature of the problem, as less firm biosynthetic assumptions can be made compared to *N*-glycans. Although automated *O*-glycan annotation approaches have been proposed to aid the determination of isomeric structures [11], the exact decisions made by such approaches are not clearly interpretable, potentially affecting transparency.

For the related area of lectin-glycan binding specificities, an approach combining rule-based machine learning with expert curation has resulted in widely used and robust guidelines for a hitherto scattered field [4]. Thus, here we present a new workflow using interpretable machine learning on a large, curated set of >237,000 *O*-glycomics spectra to derive an actionable set of rules used to identify common *O*-glycan topologies and structural isomers from tandem mass spectrometry data. We then couple the identification of diagnostic peaks with our automated fragment annotation method CandyCrumbs [11], to obtain human-understandable fragmentation events that can be used for annotation. Importantly, these rules are assessed across a wide array of experimental set-ups and analysts, resulting in (i) quantifiable rule performance, (ii) rules that are designed to work in combination with each other, and (iii) annotation confidence values of isomers identified with these rules.

Throughout this work, we also compare where our rules confirm or deviate from existing diagnostic fragments from the academic literature. We show that most *O*-glycan isomers can be confidently separated with a small number of diagnostic features, including ratios of fragment peaks, and even identify fragmentation patterns that are generalizably indicative of the same structural feature across many different glycans. We envision that this work will improve *O*-glycomics annotation accuracy, transparency, homogeneity, and accessibility, leading to new biological discoveries of the role of fine-structural details in glycans.

## Materials and methods

### Dataset construction

The herein used dataset of glycan tandem mass spectra was extended from a previously curated dataset [11]. Briefly, MS raw files were retrieved from, predominantly, GlycoPOST [12], converted into mzML format, and MS^2^ spectra were extracted into a tabular format. We then filtered our dataset to include only MS^2^ spectra of *O*-glycans (containing a reducing end GalNAc or Fuc, as well as *O*-glycan peeling products), measured in negative ion mode, and only including structures which had undergone reductive β-elimination. All annotations by experts in this dataset were assumed to be true. The final dataset consisted of 237,931 spectra and 1,647 unique glycans across 121 unique datasets (comprising 1,442 glycomics raw files).

### Data processing

Spectra were normalized by expressing their intensity as a percentage of the highest peak in the spectrum, in accordance with common practice, to facilitate direct usage of intensity threshold in obtained rules. Spectra were then binned by summing their intensities in *m/z* windows spanning 0.5 Da. Keeping track of the *m/z* difference between bin edge and peak allowed us to reconstruct the exact *m/z* later in the process [11]. Finally, we also formed relevant ratios between all bins of at least a mean value of 0.01 (i.e., 1%), as potent interaction features. Both normalized bins and ratios were available as features to the model trained to distinguish isomers.

### Decision Trees based on Shannon Entropy

In this work, we build one decision tree-based model per mass group (+/- 0.5 Da around the theoretical mass of a composition) that uses the input spectra to predict the isomers from the group. Following the divide-and-conquer idea, we do not train one decision tree for the whole problem setting; instead, we first classify the topology, if applicable, and then build separate decision trees for each topology. In early experiments, we found the performance of this approach to be superior over fitting single trees per mass group. Additionally, growing smaller trees of depths two to three was often sufficient to achieve excellent prediction performance between isomers, ensuring the practical applicability of derived rules.

Decision trees follow the idea of splitting the set of samples into two parts at each node, maximizing the purity of the partition. This means each node in a tree formulates a classification problem for the subset of samples resulting from the last splitting. The problem is solved by selecting the feature and the splitting value that best solves it. Different ways exist to measure how well a classification problem is solved. We use the Information Gain; a popular alternative is the Gini-Impurity. The Information Gain of a decision is computed as the difference in the Shannon Entropy of the node(s) above and below a split.

Figure 1A depicts how to compute the Shannon Entropy *H* as the sum over the classes *x*_*i*_ *∈ X*, with *p(x)* being the proportion of class *x* in the respective node. In this way, we can measure how pure a node is, as the presence of a few dominant classes (high *p(x)*) will lead to a low *H*. More evenly distributed class proportions result in high values for *H. H* can then be used to calculate the information gain *IG* of a split *A*, where *A* represents the splitting value of a feature, as described above. After splitting the samples based on A, the best feature and splitting value are selected by maximizing the information gain where H(X|A) is the weighted sum over the child nodes. Figure 1B visualizes that this scheme can be applied recursively until a stopping criterion is reached [13].

**Figure 1.**
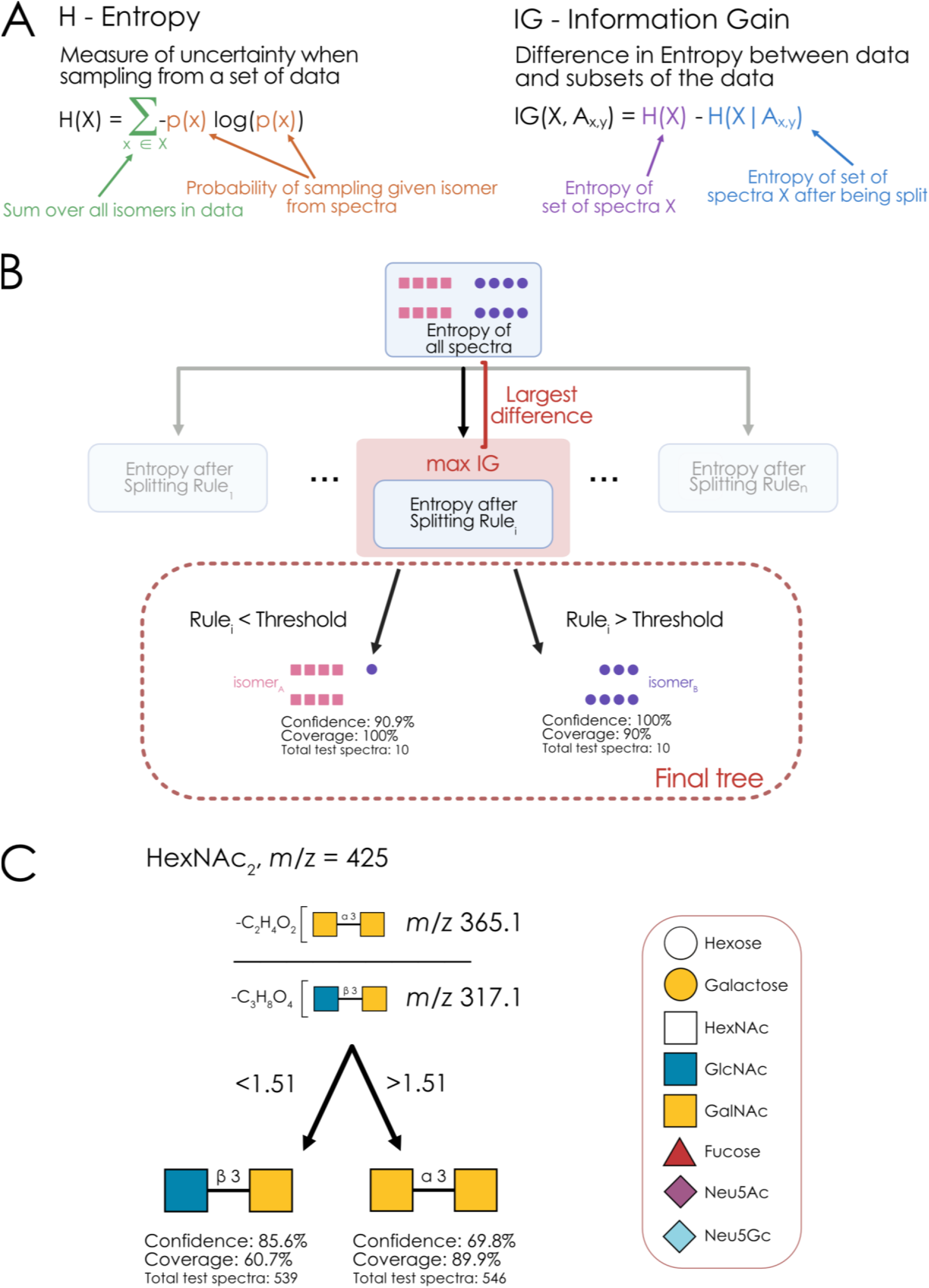
Rule-based machine learning to uncover diagnostic fragments. **A)** Definition of Entropy as a measure of sample uncertainty, as well as the Information Gain as the reduction in sample uncertainty after a given decision. **B)** Schema of decision tree construction indicating the greedy optimization of information gain at each node until the maximum depth is reached. **C)** Machine learning-derived rule for distinguishing core 3 and core 5 *O*-glycans (HexNAc_2_, *m/z* 425). The best decision tree for isomers of *m/z* 425 is shown, with the decision threshold representing values of the ratio between the two fragment ions. Confidence indicates the likelihood of a correct annotation when following the rule(s), whereas coverage designates how many spectra of that isomer follow the rule(s). The number of test spectra (not used in training the model and stemming from different experiments) for each isomer is provided in all decision trees as well. All fragments in this work are written in Domon-Costello nomenclature [20] and are visualized via GlycoDraw [16], adhering to the Symbol Nomenclature For Glycans (SNFG).

Each tree was trained using scikit-learn v1.4.2, followed by processing using glycowork v1.3 [14]. The available data per classification task was split into 70% training, 20% validation, and 10% test data. We used DataSAIL [15] for splitting to combine a similarity measure based on GlycoPOST ID and filename with stratification, to ensure each class was present in each of the splits. The trees were trained with default parameters of scikit-learn and only optimized towards their depth with the validation set. All code is available on GitHub (https://github.com/BojarLab/FragmentFactory).

### Deriving rules from trees

For each tree (both isomer and topology trees), we chose the best decision path per isomer as the source for derived annotation rules. Here, ‘best’ was determined by a score comprising the product of confidence and coverage in a leaf node for that isomer, evaluated on the independent test data (not used in any way for building the tree). Then, bins used for splitting within that decision path were mapped back to their exact *m/z* values, followed by their annotation as candidate fragments via CandyCrumbs [11], which were then visualized via GlycoDraw [16]. This resulted in a set of fragments, with corresponding decision thresholds, that could be used as annotation rules.

### Sample preparation of additional MS^n^ analyses

The sample containing HexNAc?1-?Galβ1-3(Neu5Acα2-6)GalNAc used to produce MS^3^ of the *m/z* 800 fragment and MS^4^ of the *m/z* 597 fragment was prepared from porcine gastric mucin according to the method as reported in Bechtella et al 2024 [17]. The sample containing Galβ1-3(Neu5Acα2-6)GalNAc used to produce MS^3^ of the *m/z* 597 fragment was prepared in gilthead seabream mucin as reported in Thomsson et al 2024 [18].

### Data availability

All relevant data, including their data provenance with accession IDs, can be found on Zenodo under the doi:10.5281/zenodo.12177170 [19].

### Code availability

All relevant code for this work can be found at https://github.com/BojarLab/FragmentFactory.

## Results

### Rule-based machine learning yields widely usable diagnostic fragments

A systematic approach to identify generalizable diagnostic fragments requires, at least, two things: (i) a large set of MS^2^ spectra from different experimental set-ups and different analysts, and (ii) an algorithm producing effective, but human-interpretable, rules to determine the correct isomer based on the MS^2^ spectrum. For our previous work [11], we have curated a large set of annotated MS^2^ spectra, which we have updated for this work with a special focus on *O*-glycomics data from reduced glycans in negative ion mode. Within these parameters, this dataset can be viewed as representative for a great variety of analysts and their respective set-ups. We then engaged in a rigorous data splitting procedure using DataSAIL [15] (see Methods), to ensure that we only evaluated identified rules on experiments that differed from the ones used to generate the rules. This was important to (i) ensure the generalizability of obtained annotation rules and (ii) gain accurate performance metrics (confidence and coverage) for each set of rules.

With this, we could train machine learning models to predict the annotated isomer for a spectrum, given its fragment ions. To achieve a set of annotation rules that was both performant and small, we trained a decision tree-based model for each group of isomers that minimized Shannon entropy (Fig. 1A), where each best split was considered one annotation rule (Fig. 1B). Importantly, for each isomer, this provided us with confidence and coverage values, where confidence indicated the proportion of true positives when using those rules and coverage indicated how many spectra of that isomer fell under those rules.

In general, this allowed us to construct sets of rules for many common *O*-glycan isomers where, in most cases, one or two rules were sufficient to achieve excellent confidence and coverage. One example can be seen in the model distinguishing the core 3 from the core 5 structure, where a single rule (the ratio between *m/z* 365.1 and *m/z* 317.1) was enough to effectively disambiguate between the two isomers (Fig. 1C). A value of above 1.5 here indicated the core 5 structure, allowing for easy application of this rule in practice. While there are no commonly used/accepted diagnostic fragments to distinguish these two isomers, past research comparing core 3 and core 5 structures in seabream mucin [18] supports our use of *m/z* 365, yet we here show that this can be improved by combining it with the *m/z* 317 fragment into a ratio, highlighting the potential value of this approach.

Of course, some isomeric differences, such as for the structure group Hex_1_HexNAc_1_dHex_1_ (*m/z* 530), are very robust and can be almost considered to be “solved”. In this case, the prominent presence of a HexNAc_1_dHex_1_ Z-ion (*m/z* 350.1) typically indicates an *O*-Fuc isomer (most often Galβ1-4GlcNAcβ1-3Fuc), in contrast to the standard *O*-GalNAc type isomer (Fucα1-2Galβ1-3GalNAc), in which this Z-ion would be topologically impossible. We were thus reassured to see that our new machine learning-based approach recovered these well-known effects and indeed chose *m/z* 350.1 as the best feature to distinguish these features (Fig. S1), resulting in 100% confidence and coverage at the best intensity splitting threshold. We then further aimed to distinguish type 1 and type 2 LacNAc isomers of this *O*-Fuc isomer and present *m/z* 488.2 as a potential new diagnostic fragment (Fig. S1), which indicates Galβ1-4GlcNAcβ1-3Fuc when present prominently and Galβ1-3GlcNAcβ1-3Fuc by its relative absence (given that the isomer Galβ1-?GlcNAcβ1-3Fuc has been already chosen due to *m/z* 350.1).

### Distinguishing topology and linkage differences via a divide-and-conquer approach

Many *O*-glycan structure groups comprise both topologically different isomers, as well as those differing in a single linkage, presenting a multitude of challenges to annotators. A common example of such mass groups can be found in the, still relatively modest, composition of Hex_1_HexNAc_2_dHex_1_ (*m/z* 733), which can form Lewis antigens, blood group epitopes, as well as three different core structures.

Here, we would like to showcase our divide-and-conquer approach of combining topology-level with linkage-level models to obtain effective annotation rules (Fig. 2). A fragment containing the core 3 structure (*m/z* 359) was sufficient to separate Lewis-type structures from everything else. Then, we could separate core 1 isomers of *m/z* 733 via the presence of a Y ion containing the core 1 epitope itself (*m/z* 384.2). This was then followed by the separation of blood group core 2 and core 3 structures via *m/z* 510.2, the prominent presence of which as a B-ion indicated the linear core 3 structures. Finally, a ratio of this B-ion with an A-type cross-ring fragment on the GlcNAc residue (*m/z* 409.1) was sufficient to separate type 1 and type 2 LacNAc isomers of this structure (i.e., Fucα1-2Galβ1-**3**GlcNAcβ1-3GalNAc vs Fucα1-2Galβ1-**4**GlcNAcβ1-3GalNAc).

**Figure 2.**
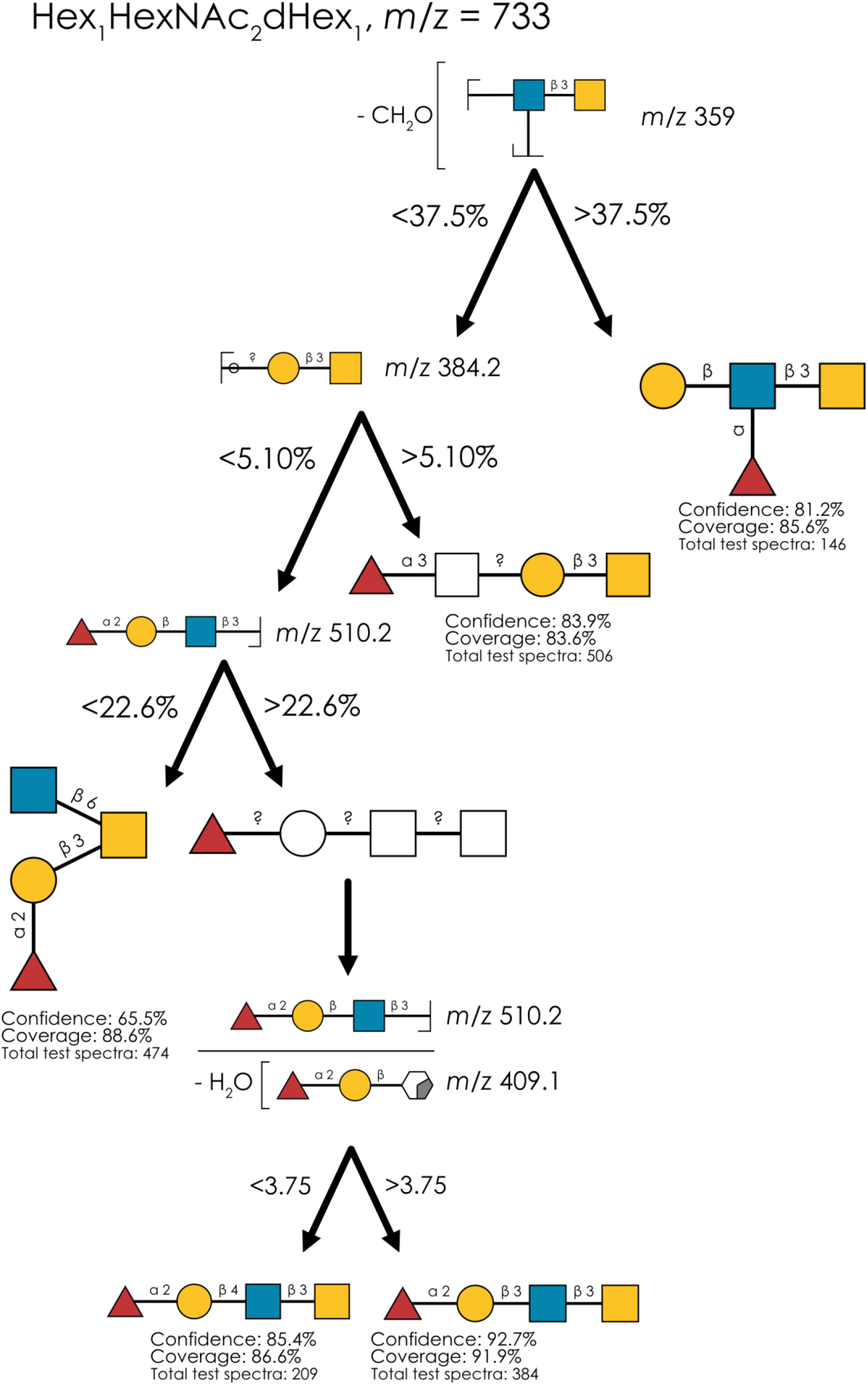
Distinguishing topologies and isomers with a divide-and-conquer approach. For the isomer group at *m/z* 733 (Hex_1_HexNAc_2_dHex_1_), we used our decision tree-based approach to find rules distinguishing topologies and, finally, isomers. The combined decision tree with all rules is shown. Rules are visualized via the SNFG-depiction of Domon-Costello fragments and their corresponding threshold values for decision making. The number of independent test spectra, as well as the therein achieved confidence and coverage, is shown for each isomer in its respective leaf node.

We were excited to see that this obtained decision scheme exhibited excellent confidence and coverage for all identified isomers. Specifically, the presented rules covered well over 80% of all spectra that contained the annotated isomers, making them extremely robust and applicable in most experimental settings. Combined with an annotation confidence of, in most cases, 80-90%, we envision these rules to raise annotation quality. We caution that, in this case, we did not identify satisfactory diagnostic features to distinguish Lewis A and Lewis X on the Lewis-type core 3 structure. The disambiguation of Lewis structures in reduced glycans presents a challenging problem in general [8, 21], which is compounded by the relative rarity of Lewis-type core 3 structures in our dataset. As discussed later, we also do want to point out that, for other mass groups such as *m/z* 895 (Hex_2_HexNAc_2_dHex_1_), our models are, in fact, capable of identifying robust indicators for Lewis A and X, respectively (Fig. S10).

### A guide to annotate common *O*-glycan isomers

Having demonstrated the capabilities of both our rule-based machine learning approach in general, as well as its extension via the divide-and-conquer approach, we then moved on to extend this potent new approach to common sets of *O*-glycan isomers. We here present a comprehensive set of quantitatively identified and characterized annotation rules for common *O*-glycan isomers (Fig. 3). We note that we only included structures in this analysis that have known and relevant isomers (e.g., no rules were constructed for sialyl-Tn antigen annotation, due to the lack of alternative isomers).

**Figure 3.**
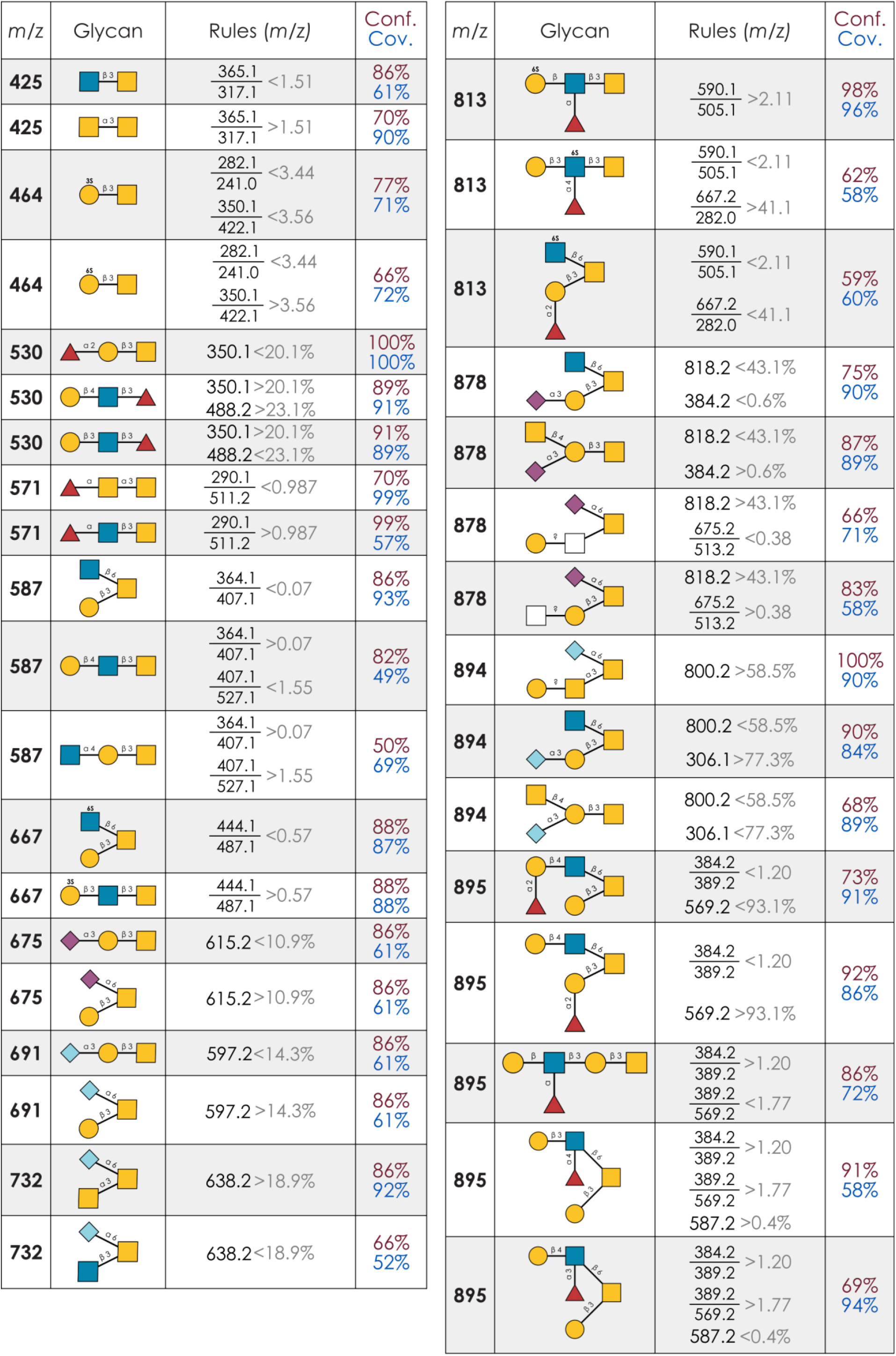
A useful guide to *O*-glycan isomer annotation. For each isomer for which we could identify performant (>60% confidence/coverage) as well as interpretable annotation rules here, we catalog the respective rules in a simplified manner. For the exact thresholds regarding intensity (individual fragments) or ratios, we refer to the respective supplementary figures (Fig. 1C, Fig. 2, Fig. 4A-D, Fig. S1-10), which list the exact models with all thresholds. Next to annotation rules, we here also depict the confidence (Conf.) and coverage (Cov.), assessed on an independent test set of experimental spectra, that result when annotating an isomer based on these rules.

Sulfated structures can be especially difficult to correctly annotate, which is why we are enthusiastic that in some cases, such as Hex_1_HexNAc_1_S_1_ (*m/z* 464; Fig. S2), our models could even identify diagnostic ratios to distinguish sulfate positioning on the galactose (Gal3S vs Gal6S) with satisfactory performance (>70% confidence and coverage). This was then extended in Hex_1_HexNAc_2_S_1_ (*m/z* 667; Fig. S5), in which we identified the ratio between *m/z* 444.1 and *m/z* 487.1 as most performant to distinguish core 2 and core 3 isomers of this composition. Other relevant examples that include new insights into diagnostic fragmentation behavior include Hex_1_HexNAc_2_dHex_1_S_1_ (*m/z* 813), a common sulfated structure group that can form either Lewis structures or an H-type 3 blood group epitope. Next to these topological distinctions, the sulfate moiety can be found on either the GlcNAc or the Gal residue, further complicating annotation. We find that a ratio of the sulfo-Lewis moiety (*m/z* 590.1) and the sulfated core 6 substructure (*m/z* 505.1) was sufficient to separate the scenarios of sulfated Gal and GlcNAc, respectively, which then was further refined via another ratio to separate Lewis and blood group structures (Fig. S7).

Overall, we note that many of the best models to distinguish isomers used ratios of fragment ion intensities as annotation features. We thus conclude, in accordance with much of the academic literature on this topic, that ratios are powerful diagnostic features and are optimistic that more complex combinations of fragment intensities, balanced with ease of use by humans, will allow for even more confident annotations. We also would like to point out that the formation of ratios is (i) more robust to systematic shifts in intensities and (ii) mitigates some of the compositional nature of relative intensities, increasing generalizability across datasets [22].

### Derived rules can generalize beyond individual structures

In general, when seeking to distinguish two specific glycan motifs or isomers, the simplest approach would be to utilize ‘topologically exclusive’ fragments, i.e., fragment masses that are only possible in a single glycan topology. Such fragments might be specific to a topology or glycan substructure, producing a high confidence value, but they are not guaranteed to occur in every experimental setup, e.g., due to preferred alternative fragmentation pathways, yielding low coverage values. To take one example, the mass of a Neu5Ac-HexNAc fragment (*m/z* 513.2) is exclusive to the topologies containing core GalNAc sialylation. This fragment has been previously described [6] as diagnostic of this type of sialylation. Yet, when tested across a more diverse set of experiments, we found it to be a rather low-coverage rule to indicate a Neu5Ac-GalNAc core motif (Fig. S8B). Specifically, the presence of *m/z* 513.2 resulted in an 86% confidence of Neu5Ac-GalNAc annotation, yet this rule only covered 57% of Neu5Ac-GalNAc containing spectra, meaning that a large fraction of Neu5Ac-GalNAc containing spectra could not be classified with such a rule.

We posit that fragments such as *m/z* 513.2 are especially preferred because they are intuitive, as they are causally related to the topology/isomer that is to be annotated. Yet, as we have shown throughout this work, annotated MS^2^ spectra contain many fragments that may not have such a clean explanation, making them less preferred for annotation, but that still offer excellent annotation quality. As a result of this, it is possible that there are many useful fragment ions not currently in use because their structure is either unknown or not intuitively thought to be connected to the isomeric difference. Our data-driven approach is designed to counteract precisely that, and we identified two such fragments that commonly occur in decision trees of sialylated structures. The fragment masses, at M-78 and M-94 for Neu5Ac/Neu5Gc, respectively, are seen in high abundance across a wide array of published MS^2^ spectra. Even when mentioned, these fragments have not been fully characterized and are either labeled simply as M-C_2_H_4_O_2_-H_2_O or, most commonly, not labeled at all.

We found that this unexplained mass loss was effective in distinguishing reducing end GalNAc sialylation from branch Gal sialylation in both Neu5Ac- and Neu5Gc-containing structures (Fig. 4A-B), though we do caution that, in an *O*-glycan context, Sia-HexNAc/Sia-Hex is conflated with α2-6 vs α2-3 linkage of the sialic acid. We can further specify this phenomenon by examining α2-6 vs α2-3 linked Sia-HexNAc motifs in milk oligosaccharides with reducing end glucose [21]. Encouragingly, both linkage types of non-reducing end Sia-HexNAc showed very low or no abundance of the M-78 fragment masses, indicating reducing end HexNAc residues are involved in this loss.

**Figure 4.**
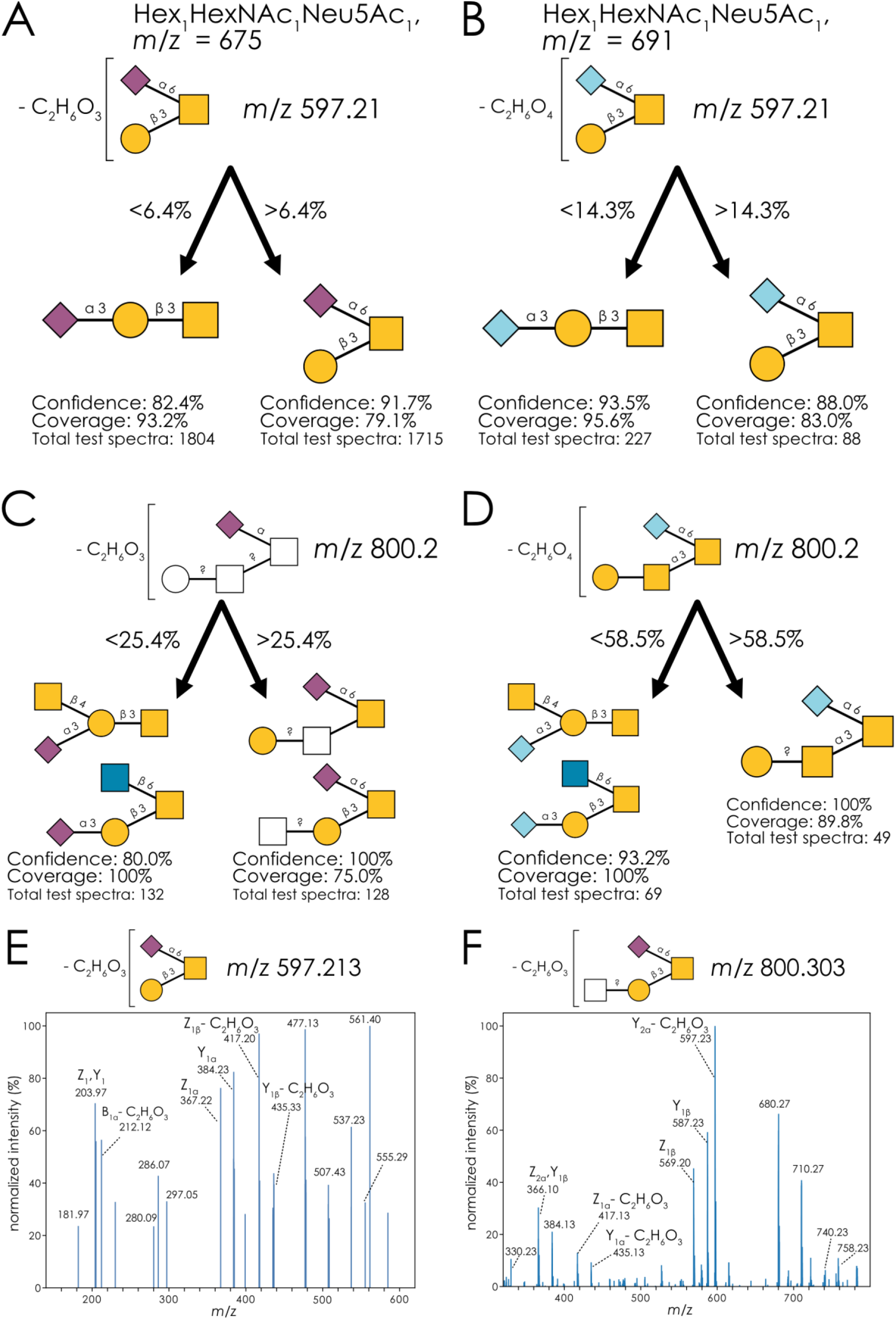
A generalizable diagnostic fragment for Sia-HexNAc annotation. **A-B)** Discriminatory performance of classifying Neu5Acα2-3Galβ1-3GalNAc and Galβ1-3(Neu5Acα2-6)GalNAc with the M-78 (C_2_H_6_O_3_) fragment (A) and Neu5Gcα2-3Galβ1-3GalNAc and Galβ1-3(Neu5Gcα2-6)GalNAc with the M-94 (C_2_H_6_O_4_) fragment (B). **C-D)** Discriminatory performance of distinguishing topologies of Neu5Ac_1_Hex_1_HexNAc_2_ with and without a sialylated reducing GalNAc with the M-78 (C_2_H_6_O_3_) fragment (C) and distinguishing topologies of Neu5Gc_1_Hex_1_HexNAc_2_ with and without a sialylated reducing GalNAc with the M-94 (C_2_H_6_O_4_) fragment (D). **E-F)** MS^3^ spectrum of the M-78 (-C_2_H_6_O_3_) fragment produced by Galβ1-3(Neu5Acα2-6)GalNAc in sea bream mucin (E) and HexNAc?1-?Galβ1-3(Neu5Acα2-6)GalNAc in porcine gastric mucin (F).

With the example of low-coverage by *m/z* 513.2 (Fig. S8B), we show that M-C_2_H_4_O_2_ (*m/z* 818.2) exhibited both higher coverage and higher confidence than the often-used *m/z* 513.2 fragment (Fig. S8C). In other work [23], this fragmentation pattern is also seen in branched sialylated trisaccharides (both Neu5Ac and Neu5Gc), as well as in larger molecules produced by extending these structures. Interestingly, Kdn-containing structures did also produce *m/z* 597 fragments, representing a loss of 36 Da (M-H_2_O-H_2_O), suggesting the losses at M-78 and M-94 to affect the C5 extension of Neu5Ac and Neu5Gc, as this moiety presents the only molecular difference. The distinguishing fragment masses in Neu5Ac and Neu5Gc differed by 16 Da, further indicating the loss to occur in the *N*-acetyl / *N*-glycolyl group of the sialic acids, due to the additional oxygen atom in Neu5Gc (Fig. 4A-B).

We thus propose that the specific fragmentation of M-78 / M-94 here presents the loss of the acetyl/glycolyl group (M-C_2_H_2_O), paired with two water losses. We note that the order of acetyl loss and then water losses was also proposed in recent work on elucidating sialic acid fragmentation in glycoproteomics data [24]. These water losses could, for instance, occur via a lactonization of the carboxyl group of C1_Neu5Ac_ with the hydroxyl group of C4_GalNAc_. Importantly, C4_GalNAc_ is axial in GalNAc, bringing the hydroxyl group into proximity of C1_Neu5Ac_, which would not be possible in the case of GlcNAc, with an equatorial C4. Using glycan 3D structure information from GlycoShape [25], we could also show that the rotational flexibility of the hydroxyl group on C4_GalNAc_ in this context was higher than that of the one on C4_Gal_ (Fig. S11), potentially explaining the diagnostic behavior of this fragmentation pattern. Another water loss, for instance via 1,7-lactonization, would then result in the observed M-C_2_H_2_O-H_2_O-H_2_O in the case of Neu5Ac-containing structures. This pattern also extended to larger structures and generalized to multiple topologies, regardless of the terminal structure on the non-sialic acid branch (Fig. 4C-D). We also note that the utility of this rule encompassed structures with an additional terminal fucose, which also yielded a high relative abundance of M-78 ions after fragmentation [23, 26].

To confirm that the losses occurred in the sialic acid moiety and not somewhere else in the glycan, we acquired an MS^3^ spectrum of this diagnostic fragment ion at *m/z* 597 (M-78; Fig. 4E). Abundant peaks at the masses representing Z_1β_-C_2_H_6_O_3_ and Y_1β_-C_2_H_6_O_3_ indicated that none of the indicated losses occurred on the galactose residue in the Hex_1_HexNAc_1_Neu5Ac_1_ isomer. Further, a substantial abundance at *m/z* 212.1 represented the commonly seen B_1α_ fragment at *m/z* 290.1, with a further loss of C_2_H_6_O_3_. Finally, the presence of unmodified Y_1α_ and Z_1α_, corresponding to sialic acid loss, support the finding that the fragmentation events of the -C_2_H_6_O_3_ loss occur only within the sialic acid. To ensure the sialic acid fragmentation was not specific to this specific trisaccharide, we acquired a separate MS^3^ spectrum of the same phenomenon in an extended structure, Hex_1_HexNAc_2_Neu5Ac_1_ at *m/z* 800 (M-78; Fig. 4F). The most abundant peak, at *m/z* 597, represented the exact same fragment ion we originally found in Hex_1_HexNAc_1_Neu5Ac_1_, which was confirmed by MS^4^ (Fig. S12). There, we identified both simple sialic acid losses at their canonical masses (Z_1β_ and Y_1β_), along with the modified losses of Galβ1-3 arm (Y_1α_-C_2_H_6_O_3_ and Z_1α_-C_2_H_6_O_3_), confirming a similar fragmentation pattern across different structures sharing this motif. While such a triple loss event would not commonly be viewed as the most parsimonious annotation explanation, we here show that it is extremely potent (high confidence), common (high coverage), and generalizable (different structural contexts), underscoring the importance of a data-driven approach to identifying diagnostic fragments in glycomics annotation.

## Discussion

Here, we presented a comprehensive resource of quantifiably performant and human-actionable rules for *O*-glycan isomer annotation based on interpretable machine learning. One of the main strengths of this work is that our annotation rules have been derived from a dataset composed of many experimental set-ups and analysts, who used different equipment (i.e., mass analyzers, collision energy, collision gas, etc.), and whose samples were present in different biological contexts, with different coeluting solutes and different solvents. Since these rules were then also validated and tested on such a diverse dataset, we can be confident that they present a more robust/performant foundation for annotation. We emphasize that our focus on coverage, typically the most neglected metric in identifying diagnostic fragments, ensures the generalizability and utility of our presented annotation rules. In principle, this process could then even be synergistically extended further, such as with retention time libraries for isomers [27], if a specific liquid chromatography context is constant for an analyst.

We are also optimistic about the promise of the herein presented workflow for further applications. In principle, the exact same workflow can be applied to the identification of similar diagnostic fragments or features for *N*-glycans, glycosphingolipids, or milk oligosaccharides. At least for some of those, the curated full dataset [11] could even be used as a data source, providing a clear and actionable implementation path. Similarly, due to the flexibility of our algorithms and CandyCrumbs [11], even data collected in, e.g., positive ion mode can be analyzed with this workflow.

We are especially enthusiastic about future work identifying further generalizable diagnostic fragments for biologically relevant motifs, similar to our efforts with Sia-HexNAc here. One example here can be found with Lewis structures, such as Lewis A and X, which currently are often only distinguished by separately analyzing non-reduced glycans [21], due to the reliance on reducing end cross-ring fragmentation as diagnostic fragments.

We caution that the herein identified rules for isomer annotation are restricted to negative ion mode and, likely, reduced glycans. As mentioned above, these are not restrictions of the workflow per se but rather restrictions of the scope that we set out for this article and, hence, stem from the used dataset. A limitation partly arising from the workflow is the possibility of additional isomers that were not considered in this analysis. A classic example could be the analysis of non-mammalian glycans [28], which may exhibit different isomers than the ones considered here, which then invalidates the use of some of the herein presented rules. We thus would like to state that the rules identified here assume that the isomers in a given tree are the only isomers that are present in major abundances in a given sample. We also advise special caution if values for ratios or individual fragments are very close to the cut-off values provided by the rules, as error rates are expected to decrease with the distance to these cut-off values.

As stated above, the preferred fragmentation pathway (ignoring collision energy as a modulator) is a function of glycan 3D structure, which then allows for the existence of diagnostic fragments to distinguish isomers in the first place. Hence, analyzing the 3D structure of isomers via molecular dynamics simulation could provide mechanistic explanations for diagnostic fragmentation, such as we have shown in previous work for distinguishing HexNAc_2_Neu5Ac_1_ / HexNAc_2_Neu5Gc_1_ isomers [11]. We envision that understanding these processes mechanistically then holds the potential of identifying more general diagnostic fragments that generalize across sequences. We are convinced there still is a need for such fragments, especially when their performance is quantified such as here, which provides (i) a standardized set of annotation rules that (ii) attaches a confidence value to annotations and (iii) overall improves the quality of annotation, leading to a more robust foundation for engaging in biological exploration of *O*-glycomics data.

## Acknowledgments

This work was funded by a Branco Weiss Fellowship – Society in Science awarded to D.B., by the Knut and Alice Wallenberg Foundation, and the University of Gothenburg, Sweden. We thank SciLifeLab and BioMS (Swedish research council) for providing financial support to the Proteomics Core Facility, Sahlgrenska Academy.

## References

1. Varki A (2017) Biological roles of glycans. Glycobiology 27:3–49. 10.1093/glycob/cww086

2. McMahon CM, Isabella CR, Windsor IW, Kosma P, Raines RT, Kiessling LL (2020) Stereoelectronic Effects Impact Glycan Recognition. J Am Chem Soc 142:2386–2395. 10.1021/jacs.9b11699

3. Zhang Z, Shah B, Richardson J (2019) Impact of Fc N-glycan sialylation on IgG structure. mAbs 11:1381–1390. 10.1080/19420862.2019.1655377

4. Bojar D, Meche L, Meng G, Eng W, Smith DF, Cummings RD, Mahal LK (2022) A Useful Guide to Lectin Binding: Machine-Learning Directed Annotation of 57 Unique Lectin Specificities. ACS Chem Biol acschembio.1c00689. 10.1021/acschembio.1c00689

5. Ashwood C, Lin C-H, Thaysen-Andersen M, Packer NH (2018) Discrimination of Isomers of Released N- and O-Glycans Using Diagnostic Product Ions in Negative Ion PGC-LC-ESI-MS/MS. J Am Soc Mass Spectrom 29:1194–1209. 10.1007/s13361-018-1932-z

6. Everest-Dass AV, Abrahams JL, Kolarich D, Packer NH, Campbell MP (2013) Structural Feature Ions for Distinguishing N- and O-Linked Glycan Isomers by LC-ESI-IT MS/MS. J Am Soc Mass Spectrom 24:895–906. 10.1007/s13361-013-0610-4

7. Doohan RA, Hayes CA, Harhen B, Karlsson NG (2011) Negative Ion CID Fragmentation of O-linked Oligosaccharide Aldoses—Charge Induced and Charge Remote Fragmentation. J Am Soc Mass Spectrom 22:s13361-011-0102–3. 10.1007/s13361-011-0102-3

8. Karlsson NG, Schulz BL, Packer NH (2004) Structural determination of neutral O-linked oligosaccharide alditols by negative ion LC-electrospray-MS n. J Am Soc Mass Spectrom 15:659–672. 10.1016/j.jasms.2004.01.002

9. Jin C, Kenny DT, Skoog EC, Padra M, Adamczyk B, Vitizeva V, Thorell A, Venkatakrishnan V, Lindén SK, Karlsson NG (2017) Structural Diversity of Human Gastric Mucin Glycans. Molecular & Cellular Proteomics 16:743–758. 10.1074/mcp.M117.067983

10. Kawahara R, Chernykh A, Alagesan K, Bern M, Cao W, Chalkley RJ, Cheng K, Choo MS, Edwards N, Goldman R, Hoffmann M, Hu Y, Huang Y, Kim JY, Kletter D, Liquet B, Liu M, Mechref Y, Meng B, Neelamegham S, Nguyen-Khuong T, Nilsson J, Pap A, Park GW, Parker BL, Pegg CL, Penninger JM, Phung TK, Pioch M, Rapp E, Sakalli E, Sanda M, Schulz BL, Scott NE, Sofronov G, Stadlmann J, Vakhrushev SY, Woo CM, Wu H-Y, Yang P, Ying W, Zhang H, Zhang Y, Zhao J, Zaia J, Haslam SM, Palmisano G, Yoo JS, Larson G, Khoo K-H, Medzihradszky KF, Kolarich D, Packer NH, Thaysen-Andersen M (2021) Community evaluation of glycoproteomics informatics solutions reveals high-performance search strategies for serum glycopeptide analysis. Nat Methods 18:1304–1316. 10.1038/s41592-021-01309-x

11. Urban J, Jin C, Thomsson KA, Karlsson NG, Ives CM, Fadda E, Bojar D (2024) Predicting glycan structure from tandem mass spectrometry via deep learning. Nat Methods. 10.1038/s41592-024-02314-6

12. Watanabe Y, Aoki-Kinoshita KF, Ishihama Y, Okuda S (2021) GlycoPOST realizes FAIR principles for glycomics mass spectrometry data. Nucleic Acids Research 49:D1523–D1528. 10.1093/nar/gkaa1012

13. Hastie T, Tibshirani R, Friedman J (2009) The Elements of Statistical Learning. Springer New York, New York, NY

14. Thomès L, Burkholz R, Bojar D (2021) Glycowork: A Python package for glycan data science and machine learning. Glycobiology cwab067. 10.1093/glycob/cwab067

15. Joeres R, Blumenthal DB, Kalinina OV (2023) DataSAIL: Data Splitting Against Information Leakage

16. Lundstrøm J, Urban J, Thomès L, Bojar D (2023) GlycoDraw: a python implementation for generating high-quality glycan figures. Glycobiology cwad063. 10.1093/glycob/cwad063

17. Bechtella L, Chunsheng J, Fentker K, Ertürk GR, Safferthal M, Polewski Ł, Götze M, Graeber SY, Vos GM, Struwe WB, Mall MA, Mertins P, Karlsson NG, Pagel K (2024) Ion mobility-tandem mass spectrometry of mucin-type O-glycans. Nat Commun 15:2611. 10.1038/s41467-024-46825-4

18. Thomsson KA, Benktander JA, Toxqui-Rodríguez S, Piazzon MC, Lindén SK (2024) Gilthead Seabream Mucus Glycosylation is Complex, Differs between Epithelial Sites and Carries Unusual Poly N-Acetylhexosamine Motifs

19. Urban J, Joeres R, Thomès L, Thomsson KA, Bojar D (2024) Navigating the Maze of Mass Spectra: A Machine-Learning Guide to Identifying Diagnostic Ions in O-Glycan Analysis

20. Domon B, Costello CE (1988) A systematic nomenclature for carbohydrate fragmentations in FAB-MS/MS spectra of glycoconjugates. Glycoconjugate J 5:397–409. 10.1007/BF01049915

21. Jin C, Lundstrøm J, Korhonen E, Luis AS, Bojar D (2023) Breast Milk Oligosaccharides Contain Immunomodulatory Glucuronic Acid and LacdiNAc. Molecular & Cellular Proteomics 100635. 10.1016/j.mcpro.2023.100635

22. Bennett AR, Lundstrøm J, Chatterjee S, Thaysen-Andersen M, Bojar D (2024) Ratios in Disguise, Truths Arise: Glycomics Meets Compositional Data Analysis

23. Jin C, Padra JT, Sundell K, Sundh H, Karlsson NG, Lindén SK (2015) Atlantic Salmon Carries a Range of Novel O -Glycan Structures Differentially Localized on Skin and Intestinal Mucins. J Proteome Res 14:3239–3251. 10.1021/acs.jproteome.5b00232

24. Geiszler DJ, Polasky DA, Yu F, Nesvizhskii AI (2023) Detecting diagnostic features in MS/MS spectra of post-translationally modified peptides. Nat Commun 14:4132. 10.1038/s41467-023-39828-0

25. Ives CM, Singh O, D’Andrea S, Fogarty CA, Harbison AM, Satheesan A, Tropea B, Fadda E (2023) Restoring Protein Glycosylation with GlycoShape

26. Zhang T, Wang W, Wuhrer M, De Haan N (2024) Comprehensive O -Glycan Analysis by Porous Graphitized Carbon Nanoliquid Chromatography–Mass Spectrometry. Anal Chem 96:8942–8948. 10.1021/acs.analchem.3c05826

27. Abrahams JL, Campbell MP, Packer NH (2018) Building a PGC-LC-MS N-glycan retention library and elution mapping resource. Glycoconj J 35:15–29. 10.1007/s10719-017-9793-4

28. Staudacher E (2015) Mucin-Type O-Glycosylation in Invertebrates. Molecules 20:10622–10640. 10.3390/molecules200610622

