## Supplemental Figures for "Navigating the Maze of Mass Spectra: A Machine-Learning Guide to Identifying Diagnostic Ions in O-Glycan Analysis"

**Supplementary Figures**

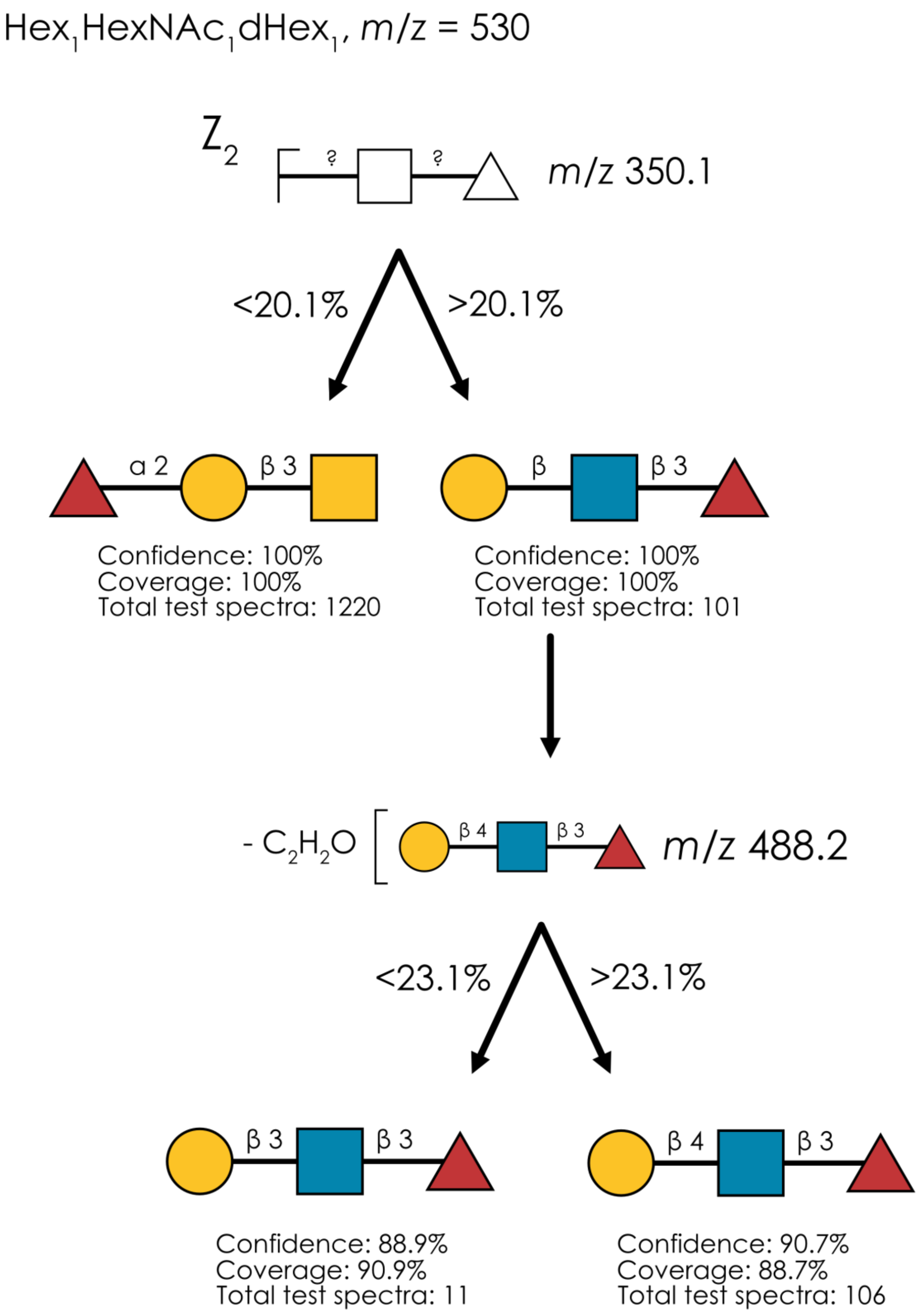


**Supplementary Figure 1. Learned annotation rules for $Hex_1HexNAc_1dHex_1$ (*m/z* 530).** Using our rule-based machine learning approach, we present the best splitting rules for distinguishing isomers of this composition. Thresholds are provided as % of the maximum intensity peak or as ratio values.

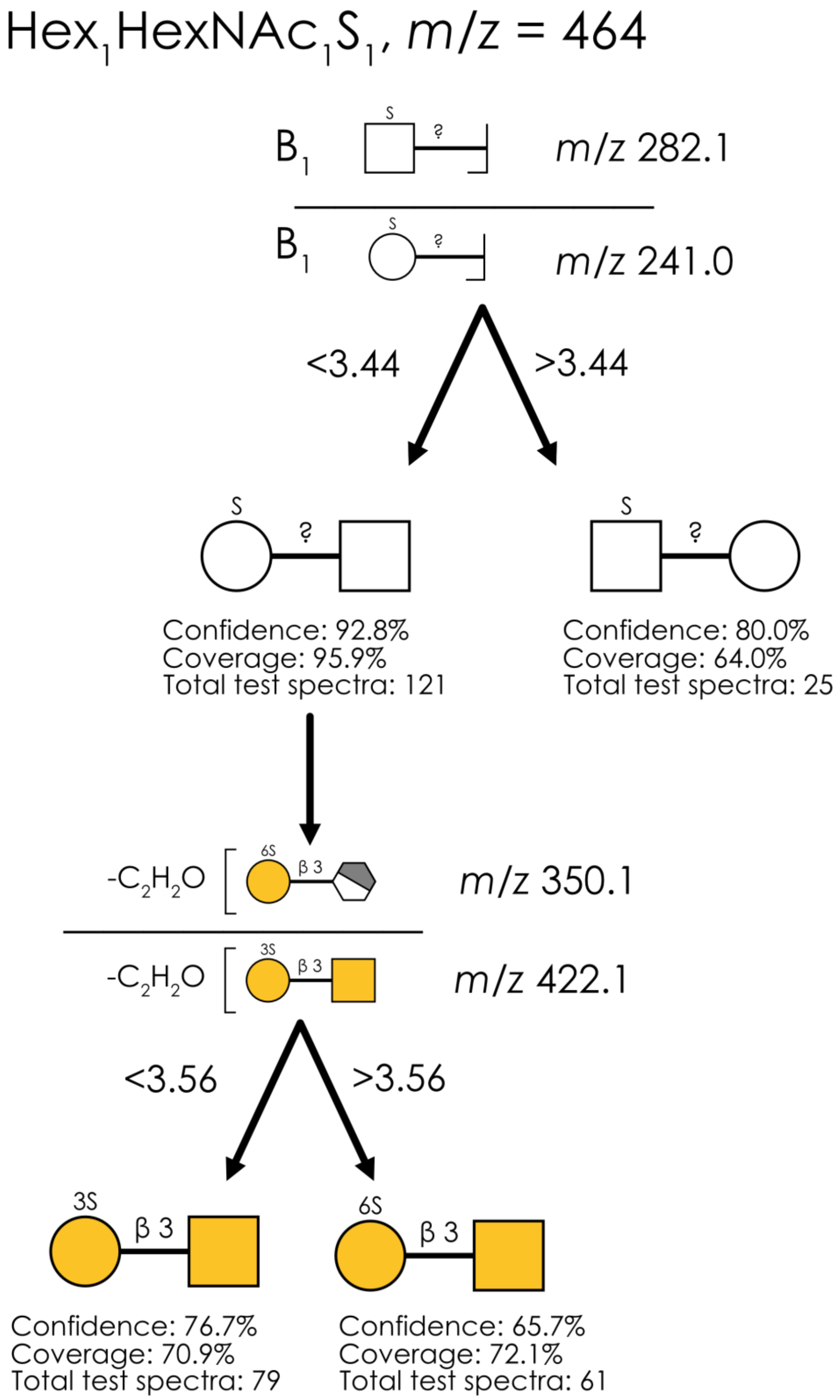


**Supplementary Figure 2. Learned annotation rules for $Hex_1HexNAc_1S_1$ (*m/z* 464).** Using our rule-based machine learning approach, we present the best splitting rules for distinguishing isomers of this composition. Thresholds are provided as % of the maximum intensity peak or as ratio values.

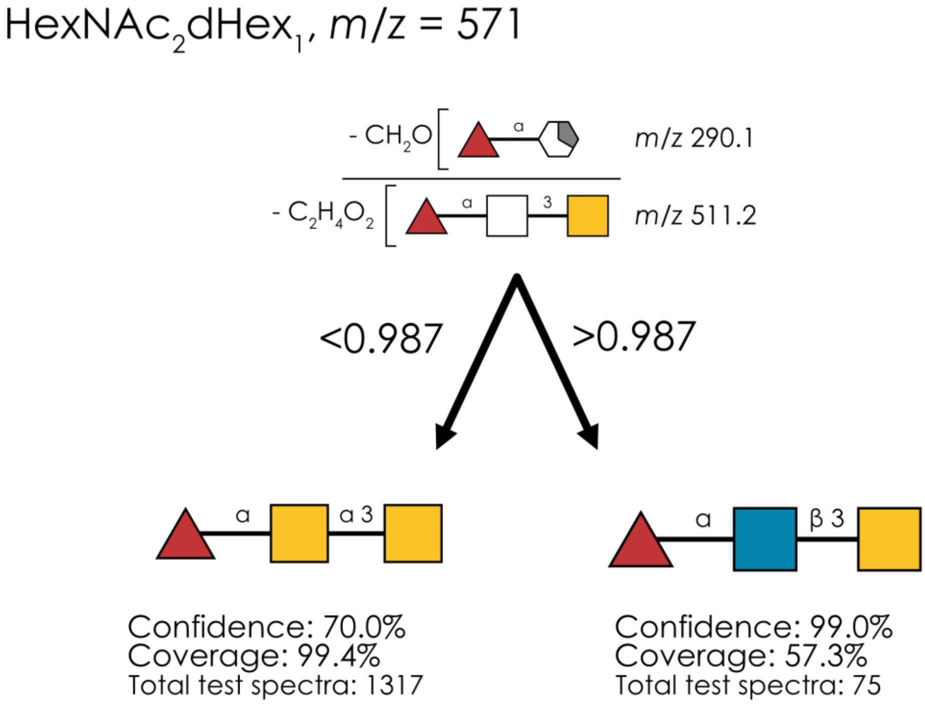


**Supplementary Figure 3. Learned annotation rules for $HexNAc_2dHex_1$ (*m/z* 571).** Using our rule-based machine learning approach, we present the best splitting rules for distinguishing isomers of this composition. Thresholds are provided as % of the maximum intensity peak or as ratio values.

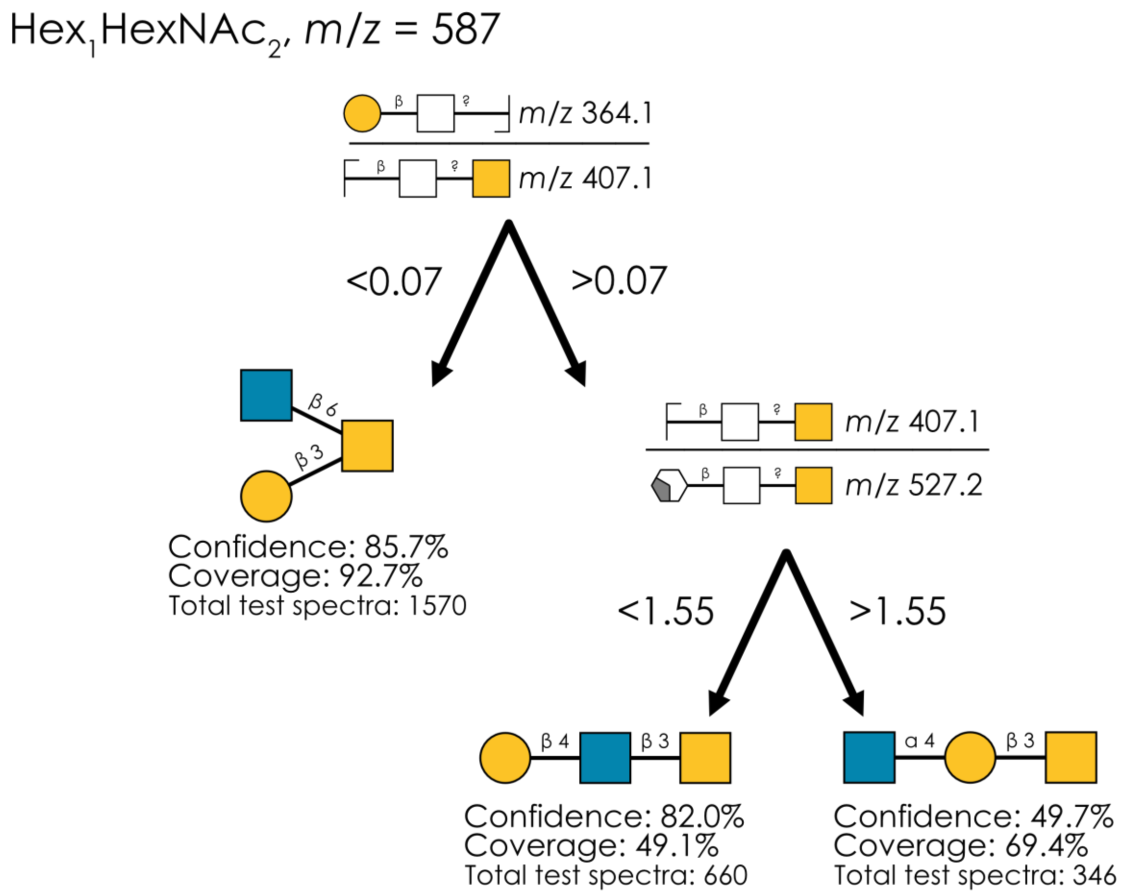


**Supplementary Figure 4. Learned annotation rules for $Hex_1HexNAc_2$ (*m/z* 587).** Using our rule-based machine learning approach, we present the best splitting rules for distinguishing isomers of this composition. Thresholds are provided as % of the maximum intensity peak or as ratio values.

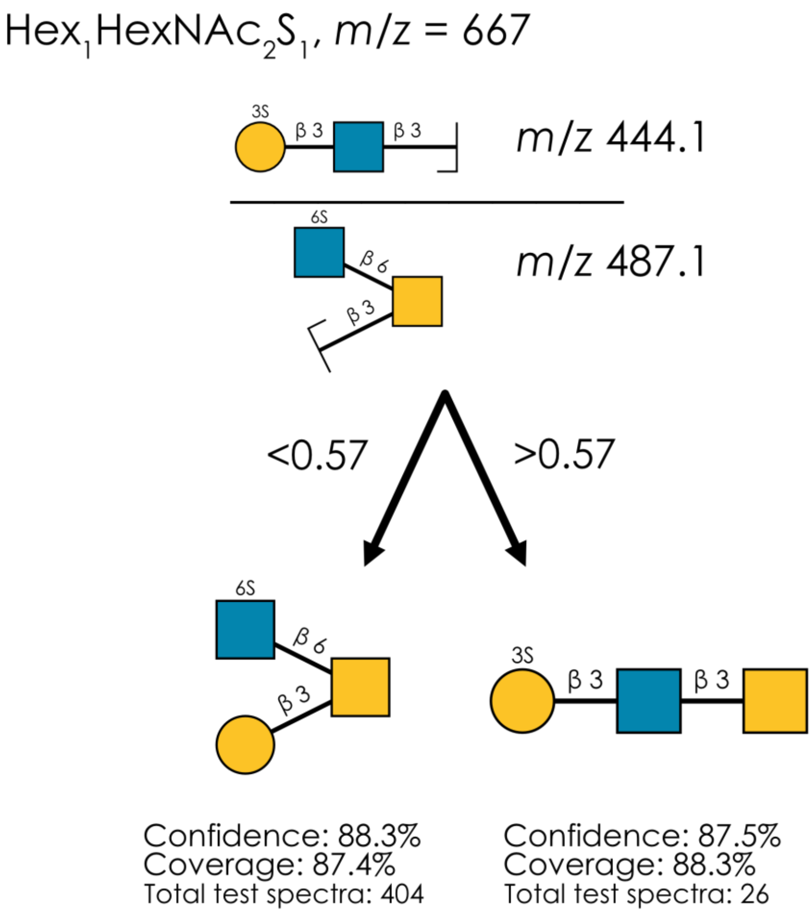


**Supplementary Figure 5. Learned annotation rules for $Hex_1HexNAc_2S_1$ (*m/z* 667).** Using our rule-based machine learning approach, we present the best splitting rules for distinguishing isomers of this composition. Thresholds are provided as % of the maximum intensity peak or as ratio values.

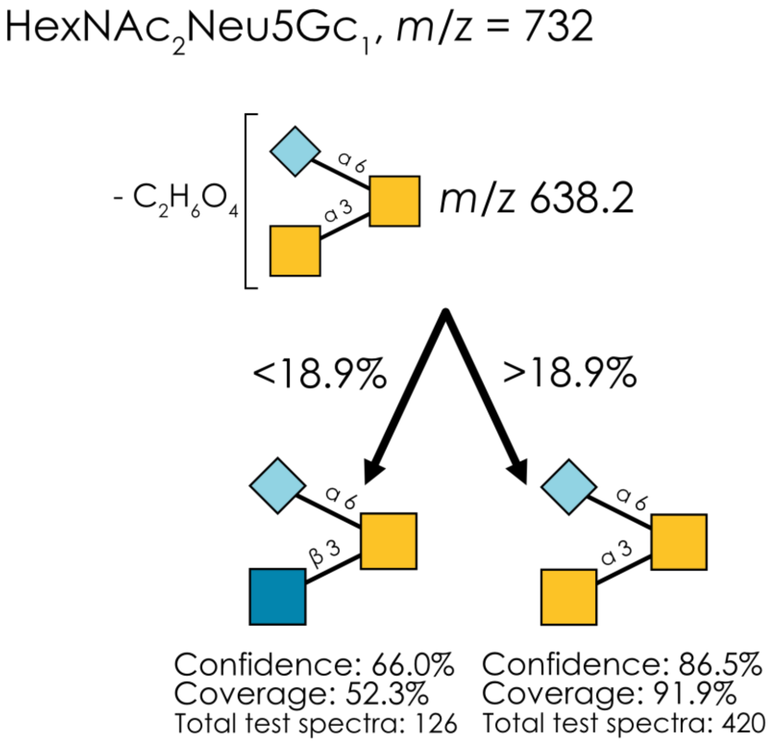


**Supplementary Figure 6. Learned annotation rules for $HexNAc_2Neu5Gc_1$ (*m/z* 732).** Using our rule-based machine learning approach, we present the best splitting rules for distinguishing isomers of this composition. Thresholds are provided as % of the maximum intensity peak or as ratio values.

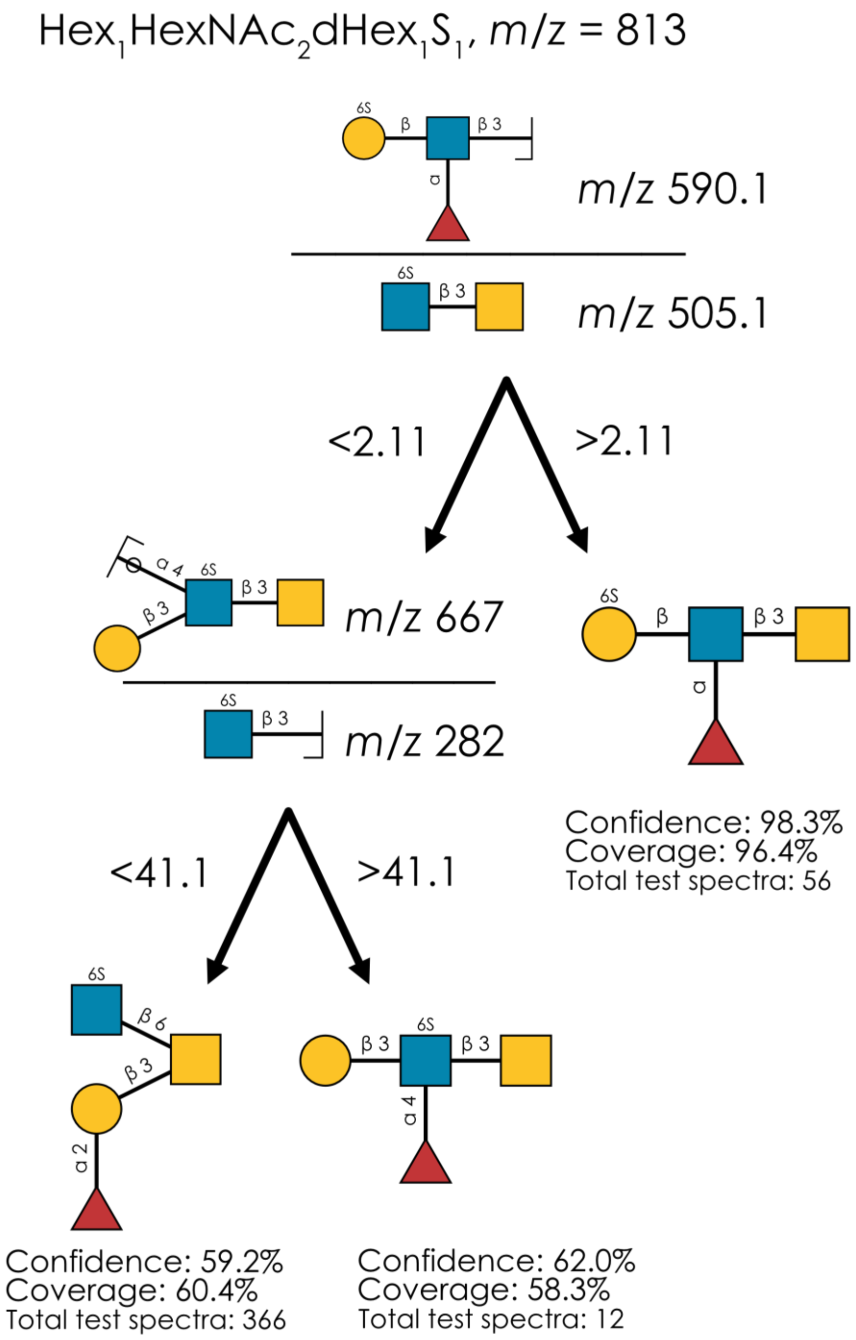


**Supplementary Figure 7. Learned annotation rules for $Hex_1HexNAc_2dHex_1S_1$ (*m/z* 813).** Using our rule-based machine learning approach, we present the best splitting rules for distinguishing isomers of this composition. Thresholds are provided as % of the maximum intensity peak or as ratio values.

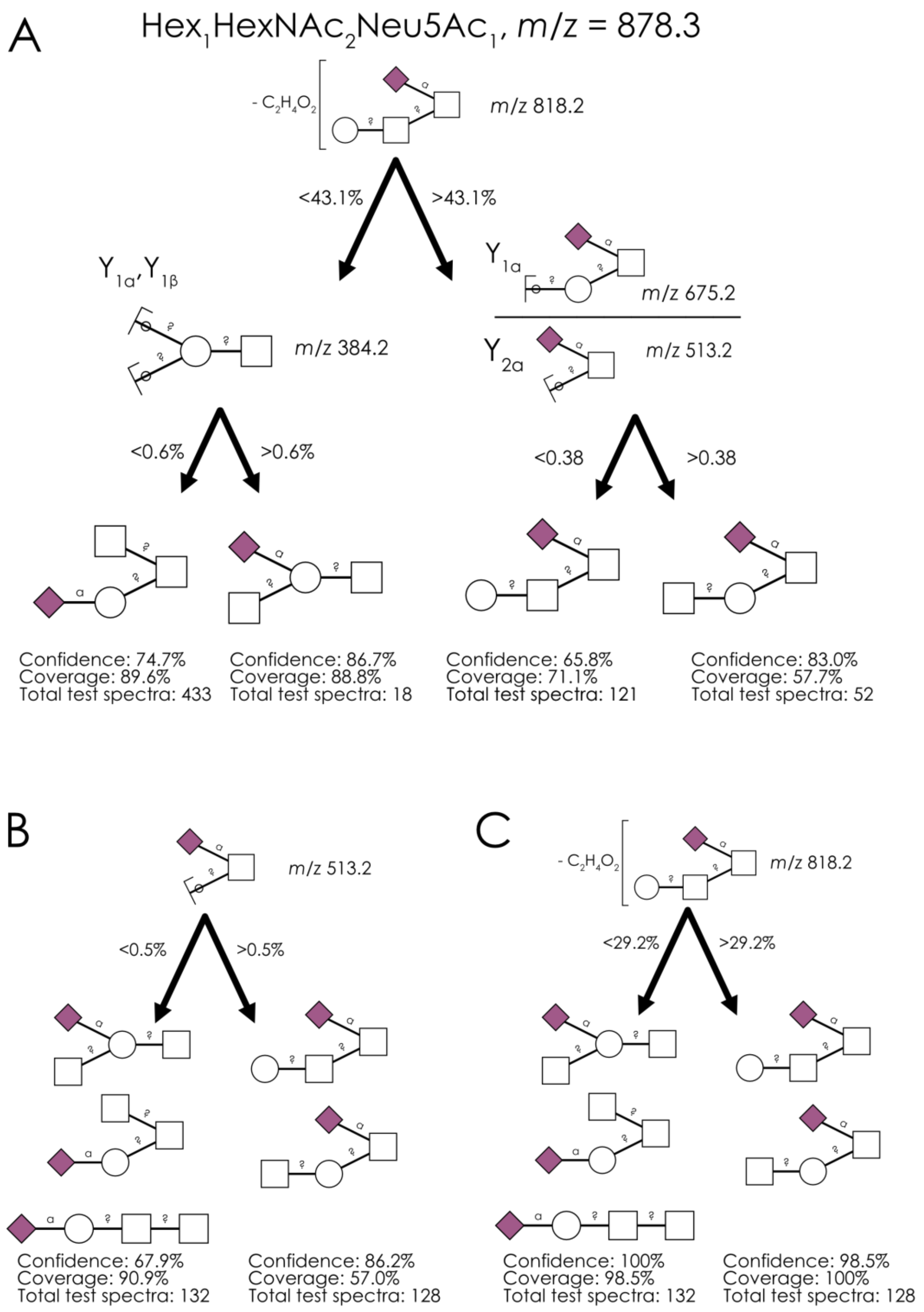


**Supplementary Figure 8. Annotating $Hex_1HexNAc_2Neu5Ac_1$ (*m/z* 878.3) with different diagnostic fragments. a-c)** Using our rule-based machine learning approach, we present the best splitting rules for distinguishing isomers of this composition. Thresholds are provided as % of the maximum intensity peak or as ratio values. The best model used M-$C_2H_2O$-$H_2O$ (*m/z* 818.2) as the topological distinguisher (a), which achieved higher confidence than using the traditional *m/z* 513.2 fragment (b). We further showed that using the M-$C_2H_2O$-$H_2O$ fragment (*m/z* 818.2), indicative of Sia-HexNAc in many other isomers, also achieved a higher average confidence and coverage than the aforementioned *m/z* 513.2 fragment (c).

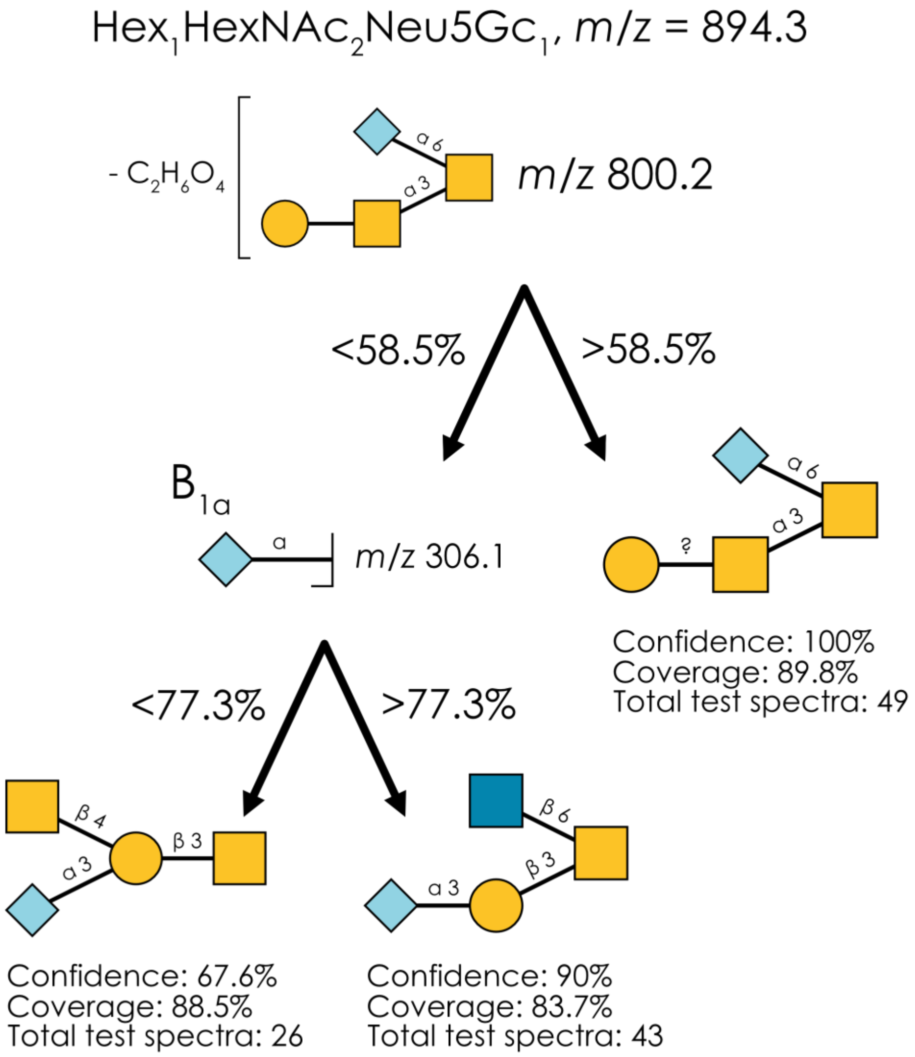


**Supplementary Figure 9. Learned annotation rules for $Hex_1HexNAc_2Neu5Gc_1$ (*m/z* 894.3).** Using our rule-based machine learning approach, we present the best splitting rules for distinguishing isomers of this composition. Thresholds are provided as % of the maximum intensity peak or as ratio values.

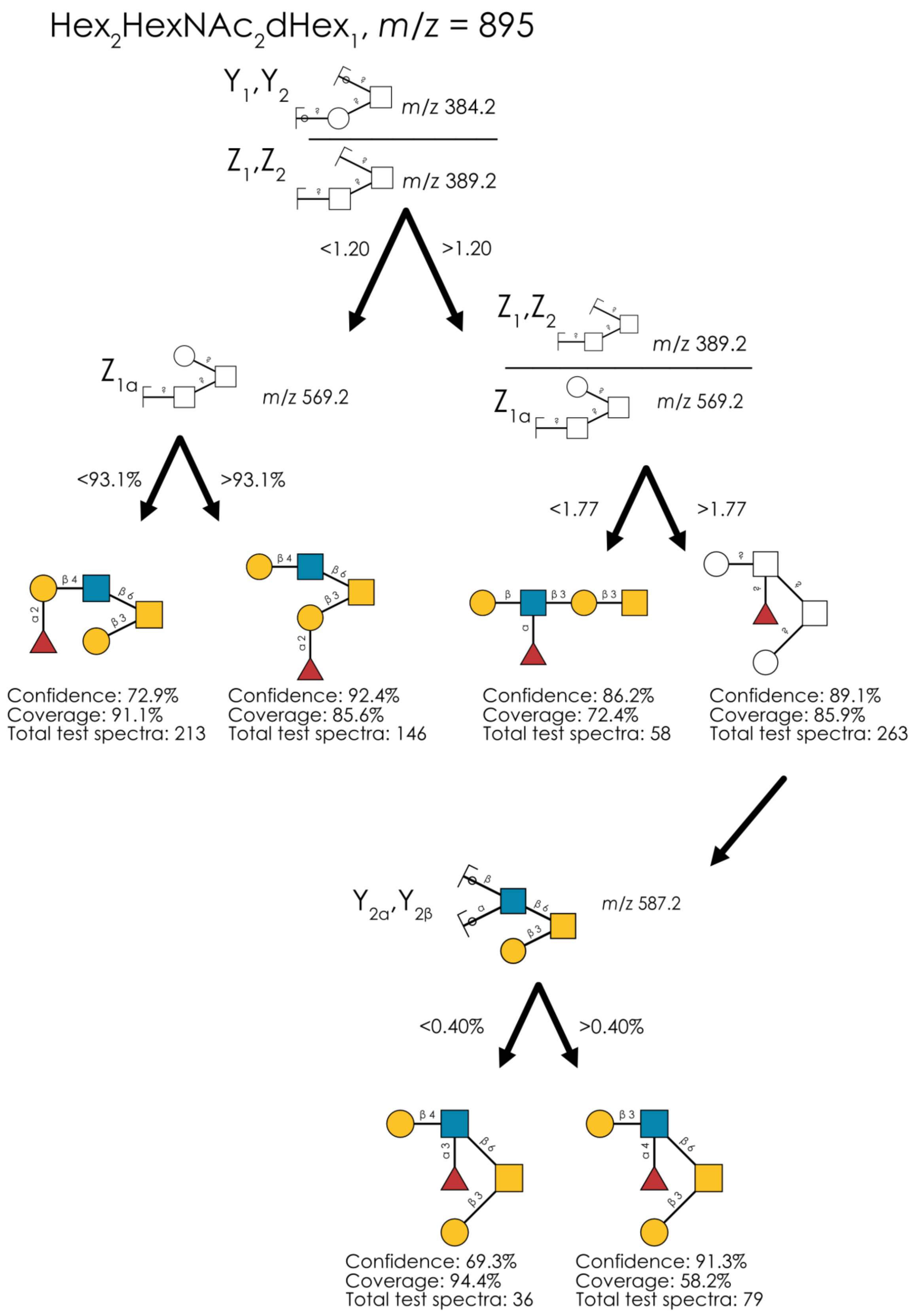


**Supplementary Figure 10. Learned annotation rules for $Hex_2HexNAc_2dHex_1$ (*m/z* 895).** Using our rule-based machine learning approach, we present the best splitting rules for distinguishing isomers of this composition. Thresholds are provided as % of the maximum intensity peak or as ratio values.

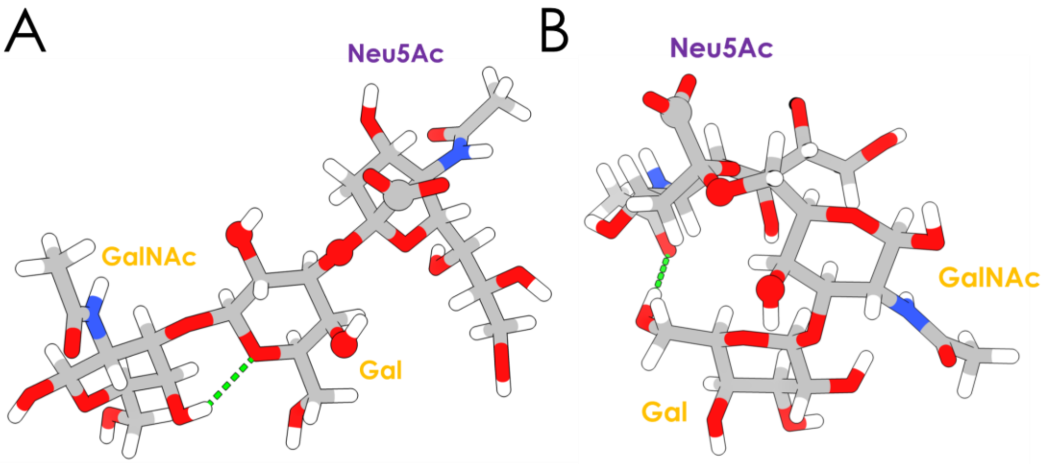


C

| Cluster N° | $Gal_{O2}$ – $Gal_{O3}$ – $Neu5Ac_{C1}$ angle (°) | $Gal_{O2}$ involved in an H-bond? | $Gal_{O4}$ – $Gal_{O3}$ – $Neu5Ac_{C1}$ angle (°) | $Gal_{O4}$ involved in an H-bond? |
|---|---|---|---|---|
| 0 | 124.8 | No | 128.7 | No |
| 1 | 129.6 | Yes | 116.4 | No |
| 2 | 166.9 | Yes | 77.4 | Yes |
| 3 | 112.1 | No | 111.5 | No |
| 4 | 106.8 | Yes | 141.3 | Yes |

**Maximal range of motion:** 60.1°

**Range of motion, excluding structurally unavailable atoms:** 12.7°

**Maximal range of motion:** 63.9°

**Range of motion, excluding structurally unavailable atoms:** 17.2°

D

| Cluster N° | $GalNAc_{O4}$ – $GalNAc_{O6}$ – $Neu5Ac_{C1}$ angle (°) | $GalNAc_{O4}$ involved in a H-bond? |
|---|---|---|
| 0 | 173.0 | No |
| 1 | 104.5 | No |
| 2 | 110.2 | Yes |
| 3 | 105.2 | No |

**Maximal range of motion:** 68.5°

**Range of motion, excluding structurally unavailable atoms:** 68.5°

**Supplementary Figure 11. Structural flexibility of $Hex_1HexNAc_1Neu5Ac_1$ isomers. A-B)** Representative 3D structures of Neu5Acα2-3Galβ1-3GalNAc (A) and Galβ1-3(Neu5Acα2-6)GalNAc (B) from GlycoShape. Green dotted lines represent predicted hydrogen bonds. $Gal_{O2}$, $Gal_{O3}$, and $Gal_{O4}$ (in A), $GalNAc_{O4}$ and $GalNAc_{O6}$ (in B) together with $Neu5Ac_{C1}$, used to compute torsion angles, are depicted with larger spheres. **C-D)** Torsion angles from all structural clusters of Neu5Acα2-3Galβ1-3GalNAc (C) and Galβ1-3(Neu5Acα2-6)GalNAc (D). Participation of $Gal_{O2}$ and $Gal_{O4}$ (C) and $GalNAc_{O4}$ (D) in predicted hydrogen bonds is also presented, impacting structural availability.

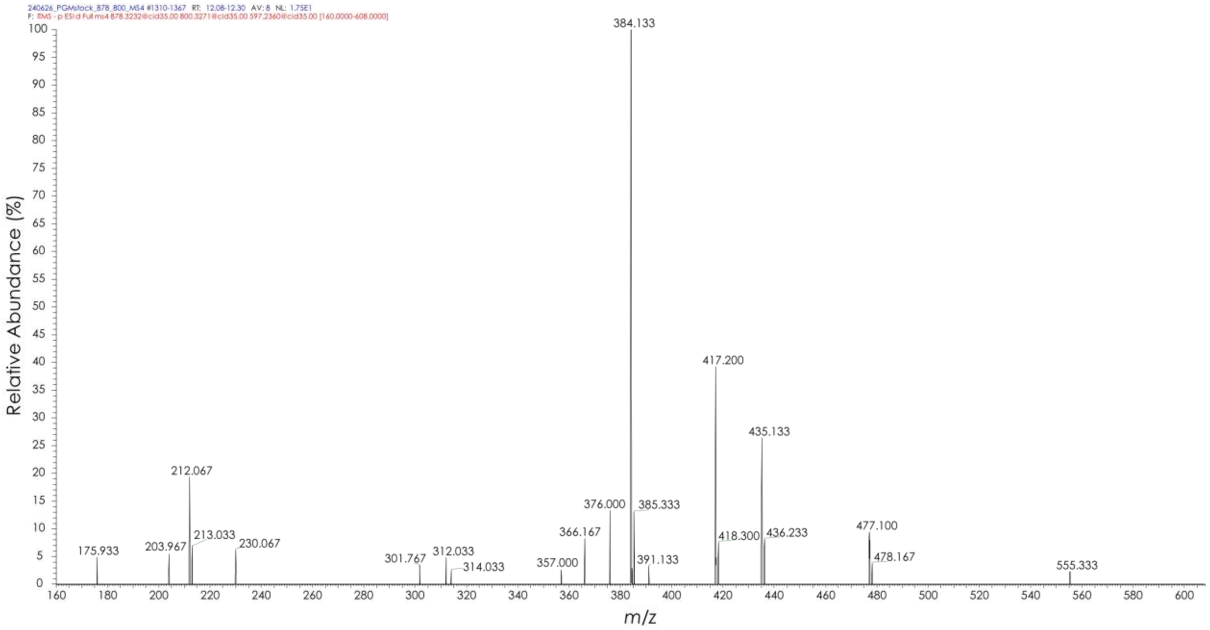


**Supplementary Figure 12. Similarity of HexNAc?1-?Galβ1-3(Neu5Acα2-6)GalNAc and Galβ1-3(Neu5Acα2-6)GalNAc diagnostic fragments.** MS$^4$ spectrum of *m/z* 597 (-$C_2H_6O_3$ -HexNAc) fragment taken from the fragmentation of the *m/z* 800 (-$C_2H_6O_3$) fragment produced by HexNAc?1-?Galβ1-3(Neu5Acα2-6)GalNAc in porcine gastric mucin.
